# The interplay of SARS-CoV-2 evolution and constraints imposed by the structure and functionality of its proteins

**DOI:** 10.1101/2020.08.10.244756

**Authors:** Lukasz Jaroszewski, Mallika Iyer, Arghavan Alisoltani, Mayya Sedova, Adam Godzik

## Abstract

Fast evolution of the SARS-CoV-2 virus provides us with unique information about the patterns of genetic changes in a single pathogen in the timescale of months. This data is used extensively to track the phylodynamic of the pandemic’s spread and its split into distinct clades. Here we show that the patterns of SARS-CoV-2 virus mutations along its genome are closely correlated with the structural features of the coded proteins. We show that the foldability of proteins’ 3D structures and conservation of their functions are the universal factors driving evolutionary selection in protein-coding genes. Insights from the analysis of mutation distribution in the context of the SARS-CoV-2 proteins’ structures and functions have practical implications including evaluating potential antigen epitopes or selection of primers for PCR-based COVID-19 tests.

## Introduction

We live in the middle of the COVID-19 pandemic caused by the Severe Acute Respiratory Syndrome Coronavirus 2 (SARS-CoV-2). In the first half of 2020 COVID-19 has spread around the world, infecting millions of people, killing hundreds of thousands, and inflicting significant economic damage. As of this writing, the future of this pandemic is still unclear. While there are signs that is it subsiding, at least in some parts of the world, it is still expanding in others and possibly coming back into areas it already affected. One of the biggest unknowns is how the pandemic would change in time. We already know that the virus is mutating and evolving into separate clades with distinct geographical and time distribution (https://www.gisaid.org) [1]. This is fairly typical for RNA viruses and it cannot be automatically interpreted as a sign that the disease it is causing is changing [2], but the first epidemiological data indicating that more transmissible viral strains are emerging [3] and preliminary experimental analyses hinting at such changes have appeared [4, 5]. The SARS-CoV-2 virus evolution is hotly debated in the scientific and the popular press [6] and has spawned a massive effort to sequence the viral variants from all over the world. A quickly growing list of complete SARS-CoV-2 genomes is maintained by resources such as GISAID [1] (https://www.gisaid.org). According to the phylogenetic analysis of this data on this site, as of early June 2020, at least six major clades: S, L, V, G, GH, and GR were identified. These clades have very uneven geographical and time distribution. For instance, most of the variants present at the beginning of the epidemic do not exist anymore; the S clade was most common in January and February 2020, while clades G, GH, and GR appeared only in February, but now constitute over 75% of new cases (https://www.gisaid.org). Over 78% of the North American isolates belong to GH (now) and S clades (early) and over 65% of isolates from the EU belong to G and GR clades. Moreover, some of these clades seem to be undergoing further splits (GISAID as of June 8^th^).

The genomic data on the SARS-CoV-2 virus is providing us with a unique understanding of the dynamics of the COVID-19 pandemic in terms of the origin of outbreaks in different locations and effects of people’s travel patterns on subsequent waves of the epidemic. For instance, the emergence of the G clade is associated with the transfer of the original infection to Italy and further to New York and the US East Coast. On the other hand, the viruses from the S clade traveled directly from China to the US West Coast and various locations in the US show a mixture of the S and G clades reflecting the pattern of human travel within the US. The COVID-19 pandemic is the first such widespread disease that is studied at such a level of molecular detail and with its growth and the analyses being updated in real-time.

The genomic data on SARS-CoV-2 is also studied for signals of positive selection in search of changes the virus may undergo while adapting to its new host (https://covid19.galaxyproject.org/evolution/). Such analyses are complicated by the combination of different types of events that influence the pandemic growth, many of which do not involve the virus’ fitness. One may naturally expect that adaptation to a new host would provide evolutionary pressure on the virus and lead to positive selection visible in the genome, but our understanding of processes leading to the emergence of new RNA viruses suggests otherwise [7]. The observed mutation pattern is most likely a result of genetic drift combined with geographical separations of localized outbreaks and the differences between the expansion of different clades are mostly due to a series of local founder’s mutations. But, at the same time, there is no doubt that the SARS-CoV-2 evolution, again typical for all RNA viruses [8], is strongly influenced by negative (purifying) selection that removes nonviable viruses [9]. Negative or purifying selection can provide information about genes’ or protein regions’ essentiality, but in general, it receives less attention than positive selection. In this manuscript, we explore the possibility that integrating information about genomic selection with that on protein three-dimensional structures would allow us to gain novel insights about SARS-CoV-2 evolution patterns.

Interestingly, negative selection was discussed in the context of protein structures at the beginning of structural biology, when Perutz and his team noticed that mutations happen rarely in the protein core [10]. These early observations were later corroborated for different organisms, classes of proteins, and evolutionary timescales (for a review see [11]). On average, the level of evolutionary conservation decreases monotonically with the increasing level of solvent exposure and is even better correlated to the measures describing density of residue packing such as Contact Number (CN) [11]. It is, however, not obvious if this trend would hold for a rapidly mutating pathogen tracked in the timescale of months, where neutral evolutionary drift is expected to be a dominant factor, and how it would behave for non-core residues indirectly important for protein function, such as protein-protein interfaces.

In parallel to the genomic characterization of the SARS-CoV-2 virus, its proteins are being characterized structurally at an equally rapid pace [12], mostly as a part of the drug discovery effort, and, as of the end of June 2020, there is direct or indirect high-quality structural information for over 60% of the SARS-CoV-2 proteome. Our group has recently developed the coronavirus3D server [13], available at https://coronavirus3D.org, to integrate information about the three-dimensional structures of SARS-CoV-2 virus proteins from the Protein Data Bank (PDB) [14] resource (http://rcsb.org) with the information on SARS-CoV-2 genomic variations retrieved from the China National Center for Bioinformation (CNCB) website (https://bigd.big.ac.cn/ncov/variation). This integration allows us to track in real-time the emergence of new trends or patterns in the evolving SARS-CoV-2 genomes in the context of the structures of its proteins.

In the first part of the manuscript, we show the evidence of the protein structure-driven mutation patterns and evaluate their frequency distribution to gain insights about types of selection pressure for individual SARS-CoV-2 proteins as well as for their functional domains and sites. In the second part of the paper we also analyze the mutation pattern of known antibody epitopes and regions used for COVID-19 diagnostic tests, showing that the continuous evolution of the SARS-CoV-2 virus can affect many aspects of the COVID-19 pandemic and that structural information on viral proteins aids our efforts to understand it.

In all the analyses we use a simple approach of assessing sequence conservation by comparing the frequency of amino acid changing mutations in a region to the background frequency in a larger region [15]. It means that, in fact, we evaluate the level of protein sequence conservation and interpret it as evidence of purifying selection. This is different from standard methods of evolutionary analysis which use the ratio of missense to synonymous mutations (dN/dS) [16]. Analyses performed directly on the amino acid level allows us to directly interpret the consequences of mutations for protein structure and function.

At the same time, we show that the variability of the frequency of missense mutations in SARS-CoV-2 genomes is much greater than the variability of synonymous mutations. Therefore, we assumed that, in the first approximation, the frequency of synonymous mutations can be treated as being constant over large portions of the genome and the frequency of missense mutations could be used as the approximate measure of the relative level of purifying selection in a given region. We mostly rely on this assumption but for longer regions and subsets of residues where it is possible, we also show the rates of synonymous mutations to demonstrate that their frequency does not significantly change between compared regions of the genome or sets of residues (thus supporting our approach of comparing the frequencies of missense mutations).

## Results

Coronaviruses have unique RNA copy-proof mechanisms [17], but despite this almost 30% of the positions (8,906 out of 29,880) along the SARS-CoV-2 genome have been mutated at least once, as can be seen by the analysis of over 24,913 high-quality genomes sequenced as of June 8^th^, 2020 on the GISAID website (https://www.gisaid.org/) (see the Methods section for the details of the protocol used to select these genomes). This percentage can only grow with further pandemic expansion and increased effort to study the viral genomes. The distribution of mutations along the SARS-CoV-2 genome has been discussed in many papers [18, 19]; here we search for new observations that could be made when incorporating information on the proteins encoded by the genome. Below we discuss several such observations based on the analysis of the variations in SARS-CoV-2 genomes obtained from GISAID as of June 8^th^, 2020. In the following, we would be analyzing distributions of distinct mutations observed at least once in the analyzed set of genomes, and not taking into account virus counts (prevalence) of mutations, as these depend on the frequency of sampling in specific geographic locations and other factors not related directly to the virus evolution. We only refer to virus counts when prevalence of mutations is particularly important in a given context e.g. when discussing the impact on epitopes or PCR-diagnostic tests.

### Distribution of mutations along SARS-CoV-2 genome and in its proteins

Mutations at a total of 8,906 genomic positions were detected in the SARS-CoV-2 genome (10,071 mutations) based on the alignment of 24,913 high-quality viral genomes. The number of genomes with mutations at any given site varies greatly and depends to a large degree on the relative extents of epidemics in different countries and level of efforts to sequence the viral genome, therefore in this section, we would not take this into account.

A large portion of mutations observed in SARS-CoV-2 genomes were missense mutations (59%), followed by synonymous mutations (34%) the rest consist of start/stop gain and loss, as well as mutations in untranslated regions (see Figure S1 for more details). When translated to the amino acid sequence, 4438 of 9926 (45%) of all amino acids in the SARS-CoV-2 proteome are mutated in at least one genome in the dataset used in this study. We plotted the distribution of missense and synonymous mutations using a moving 100 nt window along the viral genome (Figure 1A). A cluster of densely mutated regions near the 3’-terminus of the genome coincides with the boundary between ORF1ab (coding for non-structural proteins) and ORF2-ORF10 (where ORF2 codes Spike protein), coding for structural proteins. Other such regions can be mapped to the functional part of the genome as illustrated by the colored lines in Figure 1A. For instance, the region corresponding to the C-terminal domain of NSP3 was found to be significantly less mutated likely due to its key role in inducing the formation of double-membrane vesicles [20]. As seen in Figure 1B, the means and variances of missense and synonymous mutational distributions in 100 nt windows along the viral genome are significantly different (Levene-test p-value= 3.95E-18). The ratios of missense and synonymous mutations in different windows are weakly correlated (Spearman’s rank correlation coefficient R= 0.22) (see Figure 1C) implying that different mechanisms of selection could be coupled – regions under positive or negative selection are also mutated more or less often.

**Figure 1.**
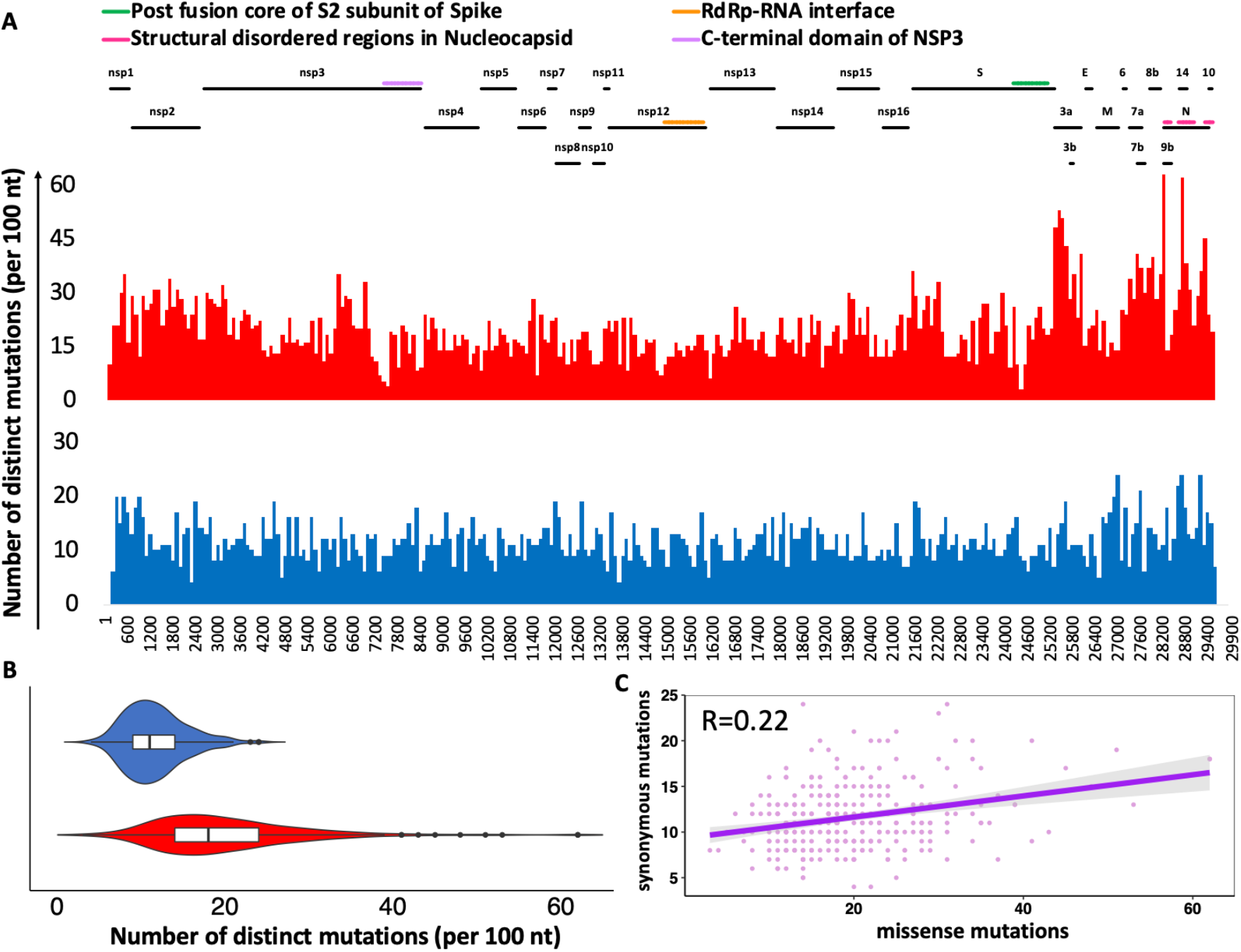
Distribution of SARS-CoV-2 SNPs based on multiple sequence alignment of 24,913 high quality genomes. A) Frequency of missense and synonymous mutations in 100 nt windows. B) Violin plots highlight the differences in the distributions of missense and synonymous mutations in windows of 100 nt. C) Correlation between distribution of missense and synonymous mutations in windows of 100 nt (Spearman’s rank correlation pf 0.22). Missense and synonymous mutations are shown in red and blue colors, respectively.

This effect is even stronger when we compare the segment of the genome for nonstructural proteins (ORF1ab, corresponding to proteins nsp1-nsp16) to that coding for “structural proteins” (ORF2-10), with the former having both a lower mutation rate and a lower ratio of missense to synonymous mutations. Both differences are statistically significant (see the details of the calculation in the Supplementary Methods). This suggests that the negative selection is stronger in the region coding for the essential virus reproduction apparatus, but also that the RNA structure of the genome supports a lower mutation rate in this region. We will use the separate mutation rates in these two regions as a background in some of the later calculations and discuss individual cases of proteins or protein regions bucking the overall trend later in the paper.

### Some SARS-CoV-2 proteins and domains show significant differences in the frequency of mutations

The SARS-CoV-2 genome codes for at least 29 individual proteins, with orf1ab product being further processed into 16 individual non-structural proteins through post-translational processing by viral proteases 3CLpro and PLpro, and some of the structural orfs coding for multiple proteins in alternative reading frames. Many of SARS-CoV-2 proteins, such as nsp3 or N protein, can be further divided into independently folding regions with specific functions. In the following analysis, we compared the observed number of missense mutations in a given protein or its region with the expected number of missense mutations under an appropriate background frequency, to identify regions that are significantly over or under mutated (see Methods). Because domain assignment is not complete for SARS-CoV-2 proteins, we use information on structurally characterized constructs to define boundaries of structural (and functional) domains or regions. Regions located in between structurally characterized domains, for instance in nsp3 protein, form another group of indirectly defined domain. The complete list of SARS-CoV-2 proteins, experimentally solved structures and domains within them that are used in the following analysis are listed in the Methods section.

In the analysis presented in this section, we focus entirely on the mutations that are manifested on the amino acid level, i.e. missense RNA mutations as they directly affect proteins, their three-dimensional structures and potentially their functions. This approach is different from the analysis and comparison of the ratios between synonymous and non-synonymous mutations that are traditionally used in the analysis of evolution of genomes. Analysis of genomic mutations on the amino acid level has a long history in the field of cancer genomics, where cancer mutations have been shown to often have clearly non-random distribution when mapped on protein sequences and structures [21]. In it worth noting that similar effects are very strong in cancer genomes where a tendency of locally increased mutation frequency is typically interpreted as a signal of positive selection and used to identify cancer driver genes and mutations.

While apparent excess of mutations in specific proteins or protein regions could be caused by positive selection, and dearth of them by negative one, this would only be true with the constant “background” mutation rate across the entire genome. With this rate undergoing significant fluctuations that appear to be random but show some dependence on the position along the genome, differences in these rates can also contribute to the excess or deficiency in the number of mutations. Therefore, in this analysis, we used different background frequencies for the different parts of the proteome being analyzed (see Methods). Table 1 presents the significant results of the analysis on individual proteins (16 nonstructural proteins and 10 structural ones) and Table 2 the results for individual functional domains and their subdomains as identified from the structural analysis. Results for both using a uniform mutation baseline and the different backgrounds for the structural and non-structural proteins are shown. The full table is available in the Supplementary Table S1. As seen from Table 1, most SARS-CoV-2 proteins do not show evidence of any selection pressure and their mutation pattern is consistent with that resulting from genomic drift. Only three proteins show statistically significant deficiency of mutations as compared to the background, specifically the spike protein (S protein), Membrane glycoprotein (M protein) and nsp12 (RNA polymerase). Nine proteins show statistically significant surplus of mutations as compared to the background.

**Table 1.** SARS-CoV-2 proteins with frequency of mutations significantly different from the background (proteome). The Bonferroni corrected significance threshold is 0.0019. “n/s” – p-values not significant even before the correction. P-values indicating significance are shown in bold font.

|  |  |  | Non-structural proteins / structural ORFs background |  | Proteome background |  |
| --- | --- | --- | --- | --- | --- | --- |
| Protein name | Length (nt) | # missense mutations | p-value greater | p-value less | p-value greater | p-value less |
| surface glycoprotein | 3822 | 762 | n/s | **1.69E-28** | n/s | n/s |
| membrane glycoprotein | 669 | 117 | n/s | **2.30E-07** | n/s | n/s |
| nsp12 | 2795 | 416 | n/s | **5.45E-06** | n/s | **2.04E-13** |
| ORF8b protein | 366 | 128 | 0.0015 | n/s | **1.59E-08** | n/s |
| nsp1 | 540 | 133 | **0.00035** | n/s | 0.02 | n/s |
| ORF14 protein | 222 | 90 | **9.69E-05** | n/s | **3.80E-09** | n/s |
| nucleocapsid phosphoprotein | 1260 | 438 | **4.01E-09** | n/s | **1.96E-25** | n/s |
| ORF3a protein | 828 | 330 | **2.50E-13** | n/s | **1.76E-28** | n/s |
| nsp2 | 1914 | 493 | **4.01E-15** | n/s | **3.18E-07** | n/s |

**Table 2.** Structurally characterized protein domains with frequency of mutations significantly different than the background (the encompassing full protein). The Bonferroni corrected significance threshold is 0.0018. “n/s” – p-values not significant even before the correction. P-values indicating significance are shown in bold font.

| Domain | Protein | Domain length | No. of missense mutations in domain | No. of missense mutations in protein | Protein length | p-value from binomial test |  |
| --- | --- | --- | --- | --- | --- | --- | --- |
|  |  |  |  |  |  | Over-mutation | Under-mutation |
| 6vyoA | Nucleocapsid | 372 | 86 | 438 | 1260 | n/s | **1.62E-06** |
| 6wjiA | Nucleocapsid | 324 | 83 | 438 | 1260 | n/s | **0.00052** |
| Region b/w 6vyoA and 6wjiA | Nucleocapsid | 249 | 119 | 438 | 1260 | **0.00011** | n/s |
| Region b/w Spike start and 6lzgB | Spike | 996 | 259 | 762 | 3822 | **7.4E-07** | n/s |
| Region b/w 2griA and 6w02A | nsp3 | 288 | 85 | 1103 | 5835 | **4.84E-05** | n/s |

From the three under mutated proteins, only one (nsp12) shows a significant result for both models of background mutations. In this case, it is possible to argue that it is the effect of purifying selection due to the functional importance of the RNA-dependent RNA polymerase (RdRp) for the SARS-CoV-2 biology. It is an interesting contrast with nsp2, which shows a statistically significant excess of mutations, again with both background mutation models. Nsp1 protein comes close, with a low, but above threshold p-value. Functions for both these proteins are unknown, but both are not directly involved in viral replication and are not essential for the in vitro reproduction. Other proteins with the relative excess of mutations (orf3a, orf8b, and orf14) are all involved in viral-host interactions. The S (spike or surface glycoprotein) and M (membrane glycoprotein) proteins show relative dearth of mutations, but only when using the structural protein-specific mutation background.

In the next step, we looked at individual domains and subdomains within SARS-CoV-2 proteins, as most of them were shown or predicted to contain multiple domains. As domains within multidomain proteins often have their independent evolutionary history and are identifiable, individual functions, differences in mutation ratios of different domains may provide a more detailed picture of their relative importance for the virus viability. We have used a similar approach in the eDriver algorithm used to identify the role of individual domains in cancer driver proteins [15]. The domain and subdomain structures of SARS-CoV-2 proteins can be identified in two steps. First, the expression of various constructs from individual proteins allowed researchers to recognize fragments that could fold independently and often can be assigned specific functions. Three-dimensional structures of many of these domains have been determined, so here we use the mapping of the SARS-CoV-2 proteins into the PDB structures/models as a proxy for domain identification (Table 2).

We also looked at regions in between the solved structures/models, assuming that these would form domains whose structures remain unsolved (Figure 2 and Table 2). In the second step, we analyzed the experimental structures and models of SARS-CoV-2 proteins to identify subdomains within them (Table 3). The complete list of domains found in SARS-CoV-2 proteins is provided in the Supplemental Material section.

**Table 3.**
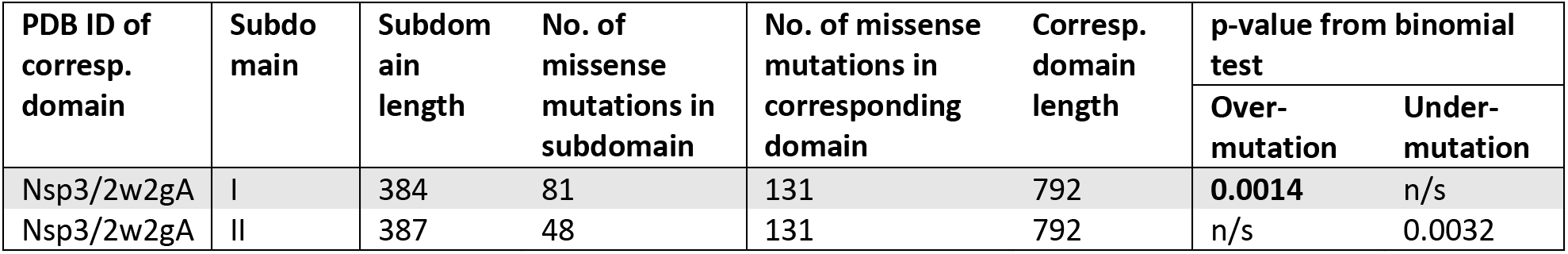
Structurally characterized protein subdomains with frequency of mutations significantly different than the background (the encompassing full protein domain – experimentally characterized structure). The Bonferroni corrected significance threshold is 0.0019 “n/s” – p-values not significant even before the correction. P-values indicating significance are shown in bold font.

**Figure 2.**
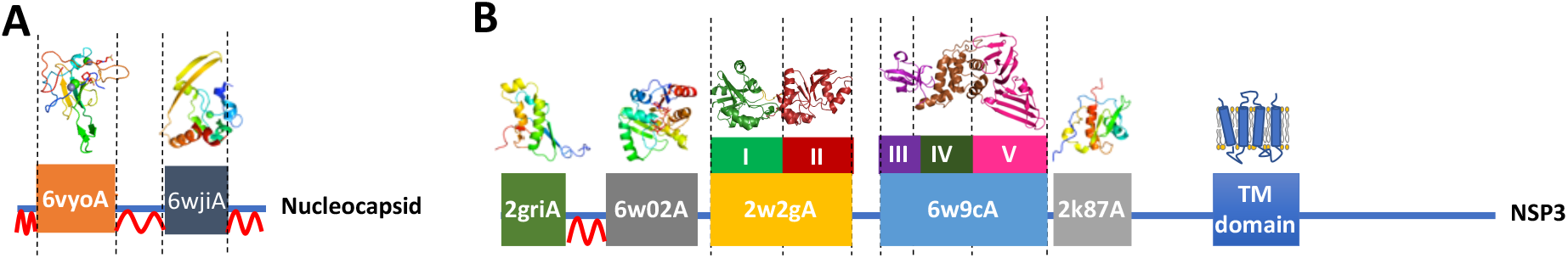
Examples of domains and subdomains used in mutation frequency analysis **A)** Nucleocapsid **B)** NSP3. Domains are labeled with PDB IDs of respective structures and models, subdomains are labeled with roman numerals, and structural disorder is marked by red wavy lines.

Full results on relative excess or dearth of mutations in all domains and subdomains are presented in Supplementary Tables S2 and S3. There are only two cases where individual domains in one protein have a statistically significant difference in mutations ratios. The structured regions in the N (nucleocapsid) protein (see Figure 2A) – the RNA binding domain and the dimerization domain (PDB IDs 6vyo and 6wji, respectively), are significantly less mutated than the remaining parts of the nucleocapsid protein. This effect could be probably explained by the rest of the nucleocapsid protein (including the region between the two domains) being partly disordered and such regions are typically much more tolerant of mutation. (for the detailed discussion of mutation rate in different parts of Nucleocapsid protein – see the section *Significantly lower mutation frequency in the region of overlapping reading frames*). The second, more interesting example is the SUD (SARS Unique Domain) region of nsp3, which consists a tandem repeat of two subdomains with a macrodomain fold and the second repeat has a significantly smaller number of mutations. Interestingly, despite strong structural similarity between the repeats, there is no obvious sequence similarity between them, suggesting that the duplication happened a relatively long time ago. These two domains cooperatively bind specific RNA structures (G-Quadruplexes) with the N- and C-terminal half playing a different role in the binding [22]. Experiments have shown that mutations in the predicted RNA biding grove in the C-terminal domain completely abolish RNA binding, while the mutations in the N-terminal domain barely affect it. This implies an essential role of the second domain and explains its relatively lower mutation rate. Interestingly, none of the residues in the predicted RNA binding site in the second domain is mutated, while all the residues in the predicted RNA binding site in the first domain were mutated at least once in the currently known genomes.

### Evidence of protein structure - driven purifying selection in SARS-CoV-2

The proportion of missense mutations in structurally characterized protein residues of SARS-CoV-2 increases with their increasing solvent exposure following known trend observed in many protein families from different organisms. There is a strong, nearly linear increase of frequency of missense mutations with synonymous mutations remaining at the approximately constant level (Figure 3A). This can be explained by tightly packed cores presenting strong constraints for amino acid residue choices and many mutations leading to unfolded protein products. Protein-protein interaction interfaces do not pack as tightly as protein cores, but also have specific amino-acid composition and their mutations may lead to function-affecting changes in protein complex formation. Notably, in cancer, we see the opposite effect, with disproportionally high number of driver mutations found on protein-protein interfaces of cancer driver genes. SARS-CoV-2 non-structural proteins are known to form higher order assemblies essential for their function[23], thus, we can expect that in most cases interface residues should be conserved. This hypothesis is confirmed by results shown in Figure 3B, where the ratio between missense and synonymous mutations fall for the residues on known protein interaction interfaces to a value between those for exposed and buried residues.

**Figure 3.**
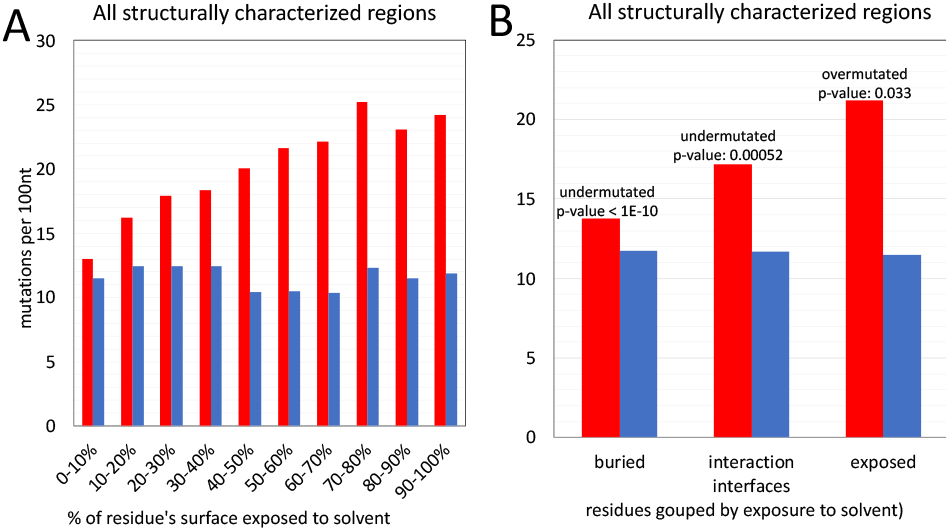
Frequencies of missense (red) and synonymous (blue) mutations for residues buried in the protein core, exposed to the solvent and involved in known protein-protein interfaces. **A)** The purifying selection decreases with increasing solvent exposure. **B)** For interfaces, the level of purifying selection falls between that for exposed and buried residues.

In the calculations shown here we only used the information on currently known protein-protein interfaces in SARS-CoV-2 proteins, based on experimental structures of viral protein complexes. We can expect that “unexplained” conserved patches on surfaces of SARS-CoV-2 proteins may aid discovery of some yet unknown interaction interfaces.

### Significantly lower mutation frequency in the region of overlapping reading frames

Overlapping reading frames are rare in eukaryotes but happen more often in bacteria and viruses, resulting in protein coding density over 100%. Systematic analyses suggest that combined negative selection on two reading frames results in decreased frequency of all mutations as mutations synonymous in one reading frame may be missense (and potentially deleterious) in another reading frame [24]. The N-terminal part of the Nucleocapsid gene of SARS-CoV-2 is translated into two different reading frames resulting in an additional gene coding for a functional protein ORF9b. A similar overlapping reading frame is suggested for the region coding for ORF14. We tested the frequency of mutations in the region of the Nucleocapsid protein which is coding for two proteins in two different reading frames and compared the mutation frequency in this region to the background frequency for the entire gene confirming the expected result (see Figure 4). However, the decrease in the frequency of mutations is only observed in the region where proteins coded in two reading frames have well-defined structures. The N-terminal region of the Nucleocapsid gene does not have an experimental structure and is predicted to be structurally disordered and, as such, is expected to impose less constraint on mutations [25]. Despite the fact that it also overlaps with ORF9b (see Figure 4), the density of mutations there is not decreased (51% of its positions have at least one mutation). Similarly, the protein-coding gene ORF14 does not show any decrease in mutation density, despite the fact that it overlaps (in a different reading frame) with the gene coding for Nucleocapsid. This, again, is probably explained by the fact that it mostly overlaps with the structurally disordered region of the Nucleocapsid protein which does not impose strong constraints on mutations.

**Figure 4.**
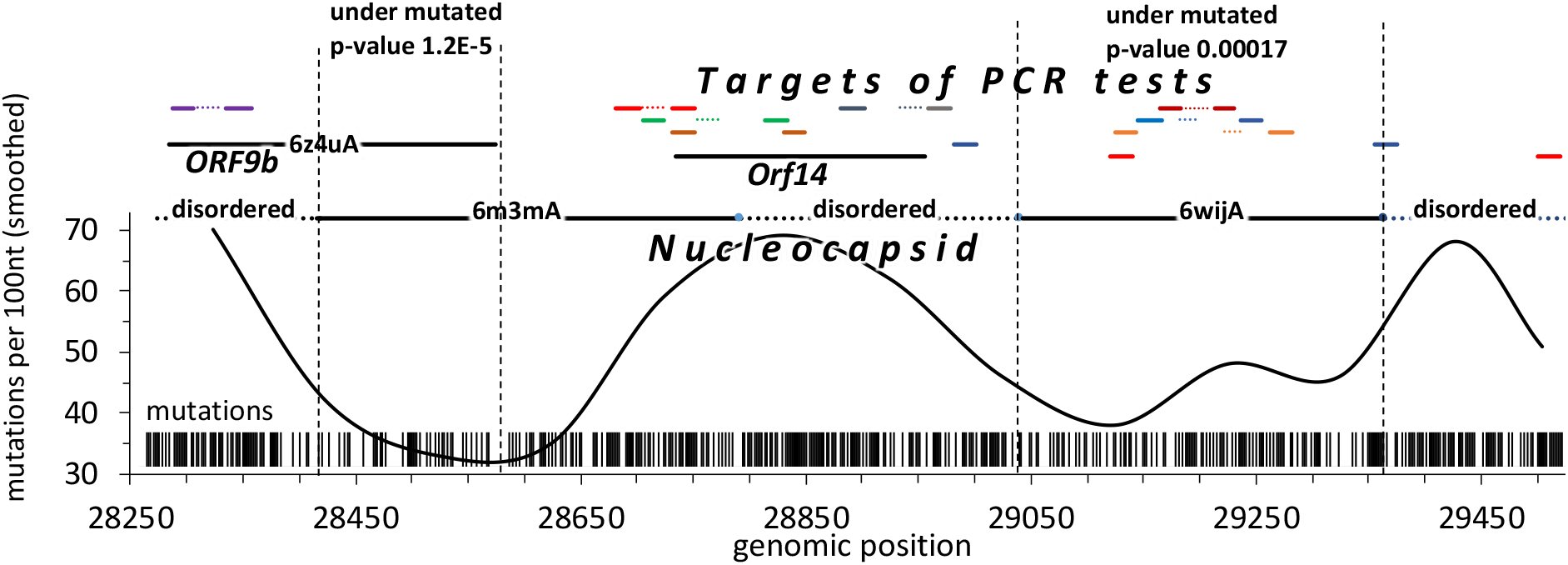
The region where structurally characterized regions of ORF9b and Nucleocapsid overlap is significantly under mutated as compared to other regions of both ORF9b and Nucleocapsid. Several target regions of PCR-based diagnostic tests for SARS-CoV-2 are in frequently mutated regions which are coding structurally disordered linkers in Nucleocapsid.

These observations have important practical implications for the selection of primers for COVID-19 diagnostic tests as mutations in their target genomic regions have detrimental effect on their accuracy. Taking into account constraints on mutation frequency imposed by structure and function conservation may help in selecting regions which are less likely to accumulate mutations in the future. In fact, multiple PCR-based diagnostic tests for COVID-19 target genome region coding Nucleocapsid protein (see Figure 4) with some of them mapping to the highly mutated disordered regions. We discuss this issue in more detail in a separate section.

### Missense mutations in known epitopes on receptor binding domain of SARS-CoV-2 Spike protein

The Spike protein of SARS-CoV-2 is the main surface antigen of SARS-CoV-2 and a preferred target of therapeutic antibodies for COVID-19 and the most likely trigger of immunological response and memory for any possible vaccine. There are already more than 15 structurally characterized complexes of various types of antibodies with Spike protein and almost all of them bind to epitopes on its Receptor Binding Domain (RBD). Mutations in the SARS-CoV-2 genome resulting in amino-acid substitutions in epitopes on the surface of Spike protein are, thus, a serious potential problem for both therapeutic antibodies and vaccines. Many of these epitopes overlap with a binding interface between RBD and ACE2 – the main entry receptor for SARS-CoV2. Protein-protein interfaces, as suggested by the results in Figure 3B, are under mutated suggesting purifying selection.

A simple comparison of missense mutation rates on the surface of Spike protein trimer, in known epitopes of antibody-Spike complexes and in buried residues reveals that the frequency of missense mutations in epitopes is slightly higher than in other surface residues (see Figure 5A). However, ACE2 interface appears to be under purifying selection with the rate of synonymous mutations exceeding the rate of missense mutations and the latter being close to that for residues from the core of the Spike protein. This is expected as the RBD interface for ACE2 is essential for the virus entry and any, even minor disruption of its binding would most likely diminish the viral ability to reproduce. As a result, the epitopes’ residues which are also involved in RBD-ACE interface are effectively “protected” from mutations and – especially – mutations which are observed multiple times (higher virus counts) – see Figures 5B and 6. In individual epitopes positions significantly involved in contact with ACE2 only rarely have mutations and these mutations usually have low viral counts (see Figure 6). Analysis of individual antibody-spike protein complexes in Figure 6 shows that even small differences in the epitopes can make a big difference, as individual residues are more prone to mutations that others. While sufficient statistics for these trends is still lacking, they support the idea that antibodies targeting epitopes with large overlap with ACE2 interaction interface might be less likely to at risk of immunological escape by the virus.

**Figure 5.**
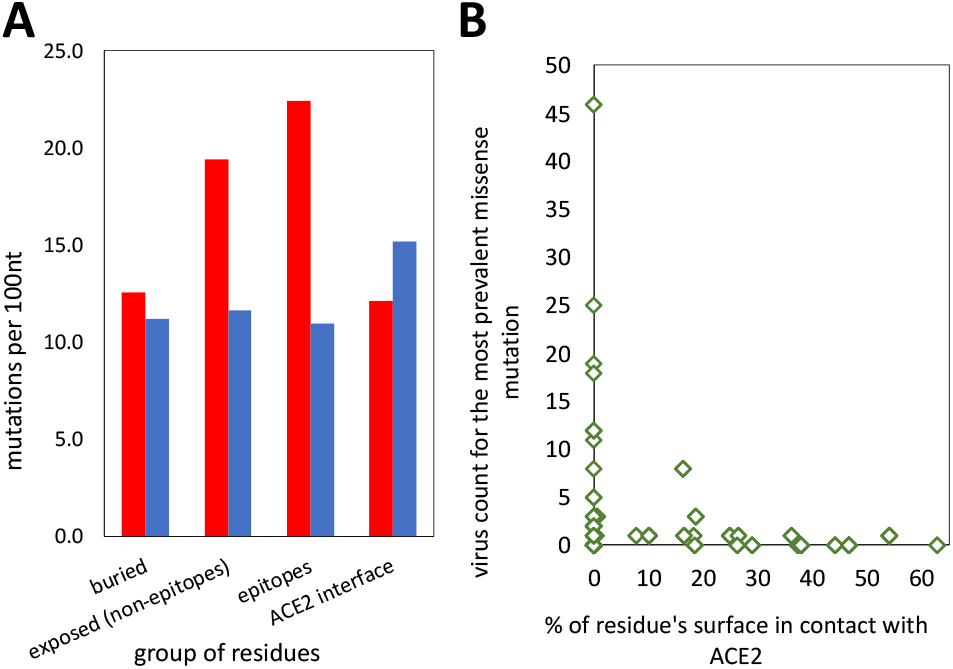
SARS-CoV-2 spike protein: **A)** Frequencies of missense (red) and synonymous (blue) mutations for positions buried in the protein core, exposed surface residues, (known) epitopes and residues involved in binding to human ACE2 receptor. **B)** The most frequent missense mutations in epitopes concentrated outside RBD-ACE interface.

**Figure 6.**
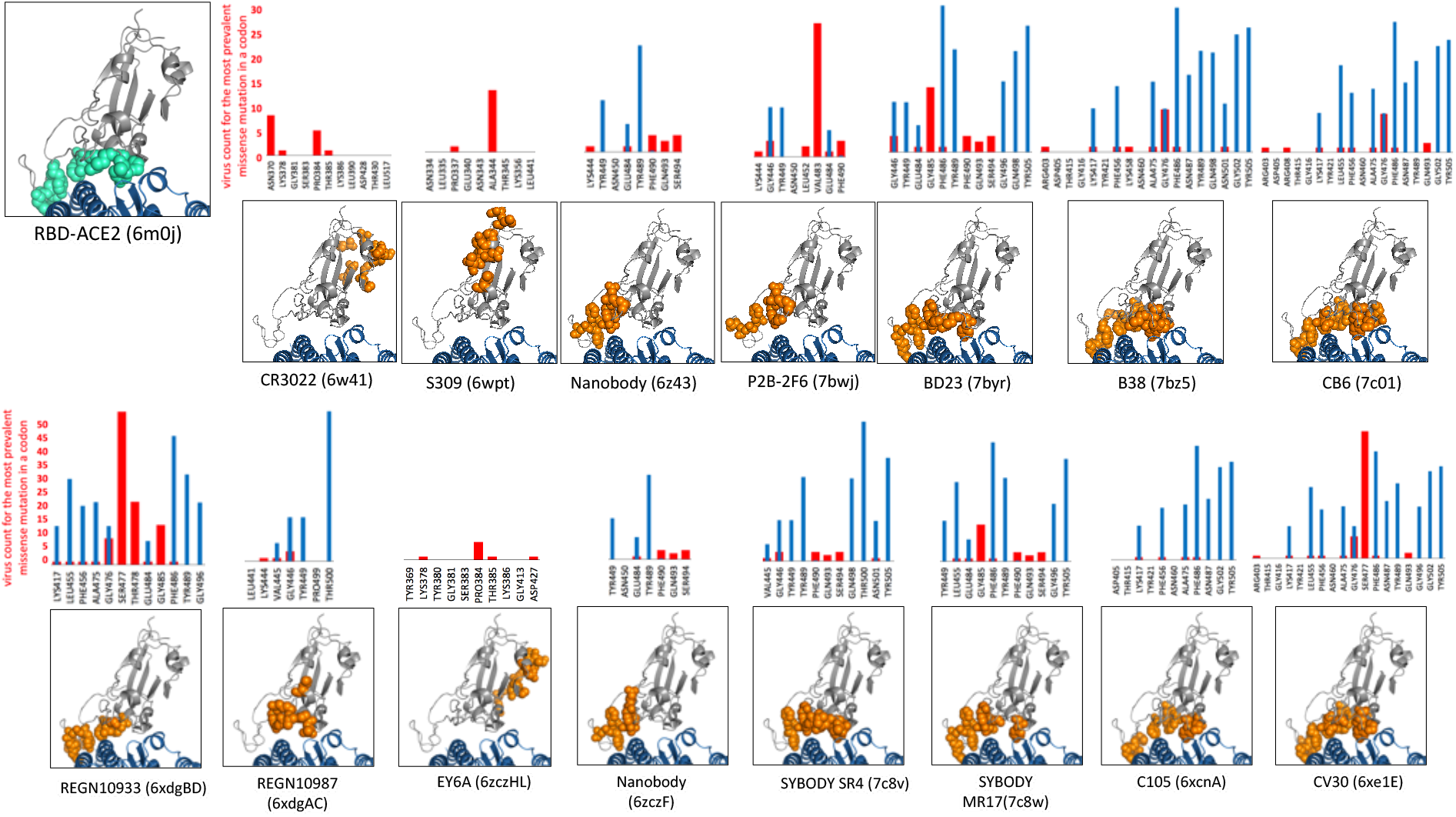
**The left-top panel** – residues involved in the interface between Spike Receptor Binding Domain (RBD – colored in gray) and ACE2 receptor – colored in aquamarine. **The 3D insets**-epitopes of antibodies binding to the RBD (RBD – gray, epitopes - orange spheres) and interface with human ACE2 receptor (blue). **The histograms** – the prevalence of missense mutations in epitopes (red bars) and % of residue’s area in contact with ACE2 (blue bars). *Epitopes* include residues whose solvent exposure decreases by at least 20% of their maximal solvent exposed area in the RBD-antibody complex as compared to RBD alone. Similarly, *ACE2-contact area* for any residue from RBD is the % of its solvent exposure lost when RBD is bound to ACE2. Antibody binding and ACE2 areas were derived from separate PDB entries (Ids of PDB entries used to define epitopes and RBD-ACE2 interface are shown above the 3D insets). For the purpose of comparison all epitopes are shown in the same structural context of the RBD-ACE2 complex (PDB id 6m0j) rather than in the context of the antibody RBD complexes. However, RBD may undergo some conformational changes in complexes with antibodies.

### The frequency of mutations in regions targeted by primers used in covid-19 PCR tests is under constraints imposed by proteins coded by these regions

The adverse effects of SARS-CoV-2 genomic mutations on PCR-based diagnostic tests potentially leading to false-negative results are widely discussed [26, 27] (also see GISAID page on popular primers available within https://www.gisaid.org/). The false-negative results of the PCR tests, especially of TaqMan-qPCR assay are linked to the high sensitivity of this technique to primer/probe-template mismatches [28] [29]. Both missense and synonymous mutations have an impact on the accuracy of PCR tests. Only the missense can be deleterious to proteins coded by mutated regions and, thus, only their frequency is linked to structural and functional constraints imposed by proteins. However, missense mutations comprise most (59%) of mutations found in the SARS-CoV-2 genome. Here we investigated the mutation rates of target regions of the widely used PCR primers and probes in relationship to proteins and protein domains coded by these regions. To this end, we collected the sequences of primers and probes commonly used for COVID-19 diagnostic PCR assays. The coordinates of genomic target regions of these primers and probes were obtained by mapping them to the reference genome used in this study (GenBank: MN908947.3) and then these genomic coordinates were mapped to SARS-CoV-2 proteins and (where possible) to experimental structures. As it could be expected, the primers targeting genomic regions coding highly conserved proteins whose functions are essential to the viral lifecycle such as RNA-dependent RNA polymerase (RdRP), show the lowest frequency of mutations (Figure 7A). In general, primer target regions corresponding to stable, experimentally verified protein structures showed lower mutation rates and prevalence (virus counts) than structurally uncharacterized and potentially unstructured regions. The structurally disordered protein regions are known to be enriched in mutations [25] and this applies to the regions targeted by some widely used diagnostic primers (Figure 7A). The example of such frequently mutated target sequences is 2019-nCoV_N1 primers and probe (also known as RX7038-N1 or CDC N1) is shown in Figure 7B. The target regions of 2019-nCoV_N1 primers/probe are coding the structurally disordered region of SARS-CoV-2 Nucleocapsid protein. Our predictions of structural disorder obtained using Disopred program [30] were recently confirmed as it was shown that the SARS-CoV-2 nucleocapsid protein is highly dynamic and constitutes of three disordered regions [31]. These structurally flexible regions are prone to mutations and are, apparently, less suitable as targets of PCR-based diagnosis of SARS-CoV-2. In contrast, the region targeted by RdRp_SARSr test has fewer mutations (Figure 7B) and, thus, seems to be a more reliable target for SARS-CoV-2 diagnostic purposes. The list of diagnostic primers and probes, mutation counts in their target regions, and proteins coded by these regions are provided in Supplementary Table S5.

**Figure 7.**
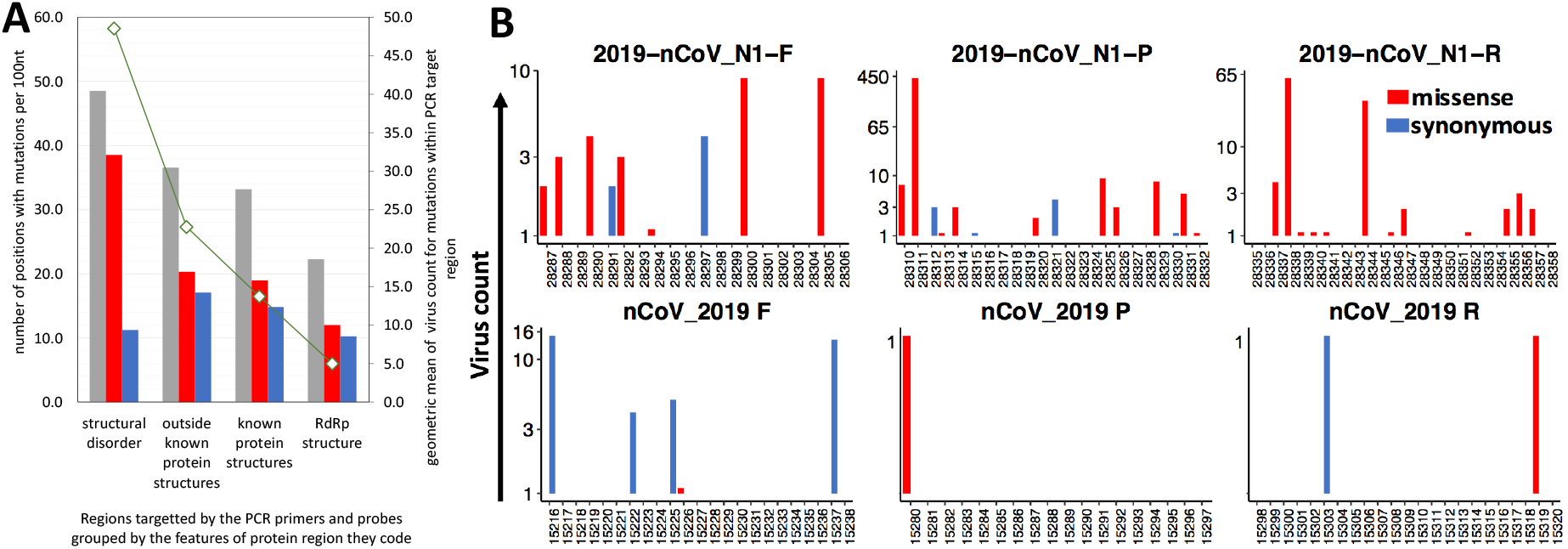
**A)** The frequency of missense mutations in regions targeted by diagnostic PCR tests is linked to constraints imposed by coded protein structures while the frequency of synonymous mutations remains roughly the same. **B)** Examples of the effects of constrains imposed by proteins on frequency of mutations in 2019-nCoV_N1 PCR test and nCoV_2019 whose target region code for disorder linker region in Nucleocapsid protein and to significantly under mutated RNA-dependent RNA polymerase (RdRP).

## Discussion

In this manuscript, we have shown that the connection between the distribution of amino acid mutations and structures of the proteins coded in the genome is already clearly evident in the genomic data currently available for the SARS-CoV-2 virus.

The rate of missense mutations significantly varies along the SARS-CoV-2 genome while the rate of synonymous mutations shows much lower variability. It indicates that mutations are significantly impacted by the selection mechanisms on the protein level.

A simple analysis of the frequency of missense mutations along the genome reveals some obvious maxima and minima. Some peaks of mutation frequency are correlated with structurally disordered regions where structural constraints of amino-acid substitution are generally lower. At the same time, some deep minima in mutation frequency correspond to known essential regions of SARS-CoV-2 proteins whose functions require high conservation of their structural details. At least one deep minimum in the rate of missense mutations corresponds to yet structurally uncharacterized C-terminal domain of nsp3 suggesting that it has a well-defined structure whose conservation is essential for the viral life cycle.

We show that nonstructural proteins have a lower mutation rate, possibly showing the effects of stronger purifying selection. It is interesting that these trends show up in the amino acid mutation ratio, but also, to a lower extent, in the rate of synonymous mutations, suggesting that some other effects on the RNA level also regulate the number of mutations.

The rate of missense mutations significantly varies between individual proteins, between functional structurally characterized domains of proteins and, even between some structural subdomains. We discuss and interpret some of these differences in the context of functions of these regions.

Mutations in SARS-CoV-2 proteins follow a known trend with positions corresponding to the residues in protein cores mutated less often that those corresponding to surface residues. Positions on the protein-protein interfaces present an intermediate case, but it is possible that the existence of some as yet unknown interaction interfaces complicate the analysis. It also opens a possibility to search for such interfaces by looking for patches of below-the-average sequence variability on protein surfaces. We are currently exploring this possibility. At the same time, disordered regions of the proteome provide the weakest purifying constraints.

Another fascinating example of constraints imposed by protein structures on SARS-CoV-2 mutations are proteins coded by overlapping reading frames. In agreement with trends observed earlier in bacteria, the region of Nucleocapsid gene coding for two different protein structures in two reading frames shows a significantly lower rate of mutations.

The analysis of the mutation pattern in the SARS-CoV-2 virus is interesting from the evolutionary point of view but maybe also of practical importance. For instance, it makes it possible to predict which of the currently known epitopes on surfaces of SARS-CoV-2 proteins are more likely to undergo widespread mutations in the future. Similar predictions can be useful for regions targeted by primers and probes used in PCR-based diagnostic tests for Covid-19, as there is already evidence that the accuracy of some of these tests is already negatively affected by the accumulation of multiple mutations [32]). We show examples from both categories, where structure constrained mutation rates may differentiate between evolutionary stable epitopes or probe sides, leading to antibodies less prone to viral escape and more reliable PCR-based diagnostic tests for COVID-19.

## Methods

### Data collection and curation

Sequences and metadata of complete SARS-CoV-2 genomes were retrieved from GISAID (https://www.gisaid.org/.) as of June 8th, 2020. All genomes containing less than 29,000 nt, low-quality genomes (i.e. those containing long stretches of NNNs) as well as genomes with high rates of mutations (above 1.5 interquartile range (IQR) from the 3^rd^ quartile mutation rate in all genomes) were removed. This filtering procedure resulted in 24,913 genomes and this set was used in all calculations and analyses in this study. One of the early annotated and sequenced complete genomes of SARS-CoV-2 (GenBank: MN908947.3) was retrieved from The National Center for Biotechnology Information (NCBI) and used as a reference for all genomic coordinates and as a query in alignments.

### Alignment, variant calling, and annotation

We calculated a multiple sequence alignment (MSA) of all high-quality SARS-CoV-2 genomes using MAFFT version 7 (https://mafft.cbrc.jp/alignment/server/) with the default parameters. The MSA file was then processed using SNP-sites[33] and BCFtools version 1.9[34] for variant calling and variant normalizations, respectively. In all analyses, we only considered single nucleotide substitutions involving unambiguous nucleotides (A,T,C,G). In the text we simply refer to them as “mutations”. All variations identified in this study along with the corresponding metadata are accessible via VarCoV application available at http://immunodb.org/varcov/.

To annotate variants, we used SnpEff (http://snpeff.sourceforge.net/). We used R package “vcfR” to manipulate and visualize variant calling format (VCF) data. The complete genome of SARS-CoV-2 (GenBank: MN908947.3) was used as a reference for genomic coordinates of proteins, protein structures, and models.

### Assessing differences in frequency of missense mutations in individual SARS-CoV-2 proteins, domains, and subdomains

For the assessment of differences in mutation frequencies of individual proteins, domains, and subdomains we compared the frequency of missense mutations in these regions to the frequency of missense mutations in some larger background region encompassing a given region of interest.

We used the binomial test to identify individual proteins, domains or subdomains that are significantly over-or under-mutated, when compared to an appropriate background mutation rate. This approach was used previously by our group in the eDriver algorithm [15] to evaluate the significance of differences in mutation rates between domains of cancer driver proteins.

The arguments for the binomial test, which are the number of successes, the number of trials, and the expected probability of success, were set as follows:

1. The number of successes was the observed number of missense mutations in the protein/domain/subdomain being analyzed. This was counted as the total number of distinct missense mutations in that protein/domain/subdomain observed in at least one sample. Therefore, virus counts (the number of samples where mutation was observed) were ignored, since we assumed that, in most cases, these would not represent independent mutation events. However, missense mutations that occurred at the same genomic position, but resulted in different base substitutions were counted independently.
2. The number of trials was the number of missense mutations in the background region used for comparison.
3. The expected probability of success (under the null hypothesis) was equal to the length of the protein/domain/subdomain divided by the length of the background region.

All lengths were calculated in terms of genomic positions (i.e., the length of the genomic region coding for the protein/domain/subdomain being analyzed). Missense mutations were also counted at the level of genomic positions.

The following approaches and background regions were used in the analyses of individual proteins, domain, and subdomains:

1. For individual proteins, we performed the binomial test in two different ways, using two different backgrounds:

a. We used the entire proteome (see note below) as the background. That is, that the number of trials was equal to the total number of missense mutations in all proteins and the total length of all proteins was used for the calculation of the expected frequency of success.
b. We used the set of all non-structural proteins (see note below) as the background for analysis of individual non-structural proteins, and the set of all structural proteins (see note below) as the background for the analysis of individual structural proteins. The full list of proteins analyzed can be found in Supplementary Table S1. Note: Two very short peptides Orf3b and NSP11 coded in alternative reading frames (containing 9 bases and 38 bases respectively) were excluded from these analyses.
2. Domains were identified based on protein structures or models. Only structures/models representing segments of the protein and not the full protein were considered. We also considered regions in between known structures/models to represent domains as well. The full list of domains can be found in Supplementary Table S2. For each domain analyzed, the encompassing full protein was used as the reference background region.
3. Subdomains were identified by analyzing known structures/models. The full list of subdomains can be found in the Supplementary Table S3. For each subdomain analyzed, the encompassing domain (structure/model) was used as the reference background region.

### Structural coverage of the SARS-Cov-2 proteome and derived structural characteristics

The structural data for biological assemblies of SARS-Cov-2 was downloaded from Coronavirus3D server developed recently by our group[13]. The Coronavirus3D server provides links to experimental structures of SARS-CoV-2 proteins stored in PDB[14] and models of protein regions of SARS-Cov-2 for which direct structural characterization is still lacking. Models were calculated with Modeller[35] based on FFAS[36] alignments. For the purpose of this study we prepared a nun-redundant list of structures which included non-overlapping structures and models providing only one structural characterization for each residue where possible (with an exception of structures coded in two different reading frames). The list of structures and models used in this study is provided in Supplementary Table S4.

The collected experimental and modeled biological assemblies of SARS-CoV-2 proteins were used to calculate solvent exposure with the GetArea program[37]. Solvent exposure was calculated separately for biological assemblies and for isolated chains. The, buried residues were defined as those with less than 20% of their surface exposed to the solved according to GetArea. The remaining residues were classified as exposed. Interfaces were defined as a subset of residues whose solvent exposure decreased by at least 20% of their total area in biological assembly as compared to an isolated chain.

### Assessment of mutation frequencies as function of solvent exposure

The list of single nucleotide mutations in SARS-CoV-2 genomes (prepared as described in the section *Collection and curation of SARS-CoV-2 variation data*) was merged with the solvent exposure data prepared for residues of SARS-CoV-2 proteins (as described in the previous section). The total numbers of synonymous and non-synonymous mutations were then calculated for codons of protein residues for different ranges and categories of solvent exposure. The significance of differences in frequency of missense mutations between buried, exposed, and interface residues was again assessed using binomial tests as described in previous sections with the entire proteome of SARS-CoV-2 used as the background.

### Frequency of mutations in overlapping reading frames

The significance of the changes in frequency of all (missense + synonymous) mutations in different regions of ORF9b and Nucleocapsid proteins was calculated using binomial tests in a way analogous to that used for individual proteins (see the previous section). For example, the number of all mutations in the region of the overlap of two structures coded in different reading frames (positions 28415-28574), the total number of mutations in Nucleocapsid and the ratio of the length of the overlap to the total length of Nucleocapsid were used as number of successes, number of trials and background probability in binomial tests, respectively. All lengths were calculated in terms of nucleotides.

## Acknowledgments

We acknowledge efforts of all the laboratories and teams responsible for obtaining the specimens, generating genetic sequence data and protein structure data and the teams at PDB, GISAID, and CNCB for maintaining and distributing this information. We thank Arash Iranzadeh for assistance in calling the SARS-CoV-2 variations. This work is sponsored in whole or in part by NIH institutes: NIAID under contract no. HHSN272201700060C and NIGMS by a grant GM118187.

## References

1. Elbe S, Buckland-Merrett G. Data, disease and diplomacy: GISAID’s innovative contribution to global health. Glob Chall. 2017;1(1):33–46. Epub 2017/01/10. doi: 10.1002/gch2.1018. PubMed PMID: 31565258; PubMed Central PMCID: PMCPMC6607375.

2. Grubaugh ND, Petrone ME, Holmes EC. We shouldn’t worry when a virus mutates during disease outbreaks. Nat Microbiol. 2020;5(4):529–30. Epub 2020/02/20. doi: 10.1038/s41564-020-0690-4. PubMed PMID: 32071422; PubMed Central PMCID: PMCPMC7095397.

3. Korber B, Fischer W, Gnanakaran S, Yoon H, Theiler J, Abfalterer W, et al. Spike mutation pipeline reveals the emergence of a more transmissible form of SARS-CoV-2. bioRxiv. 2020.

4. Lorenzo-Redondo R, Nam HH, Roberts SC, Simons LM, Jennings LJ, Qi C, et al. A Unique Clade of SARS-CoV-2 Viruses is Associated with Lower Viral Loads in Patient Upper Airways. medRxiv. 2020.

5. Daniloski Z, Guo X, Sanjana NE. The D614G mutation in SARS-CoV-2 Spike increases transduction of multiple human cell types. bioRxiv. 2020:2020.06.14.151357. doi: 10.1101/2020.06.14.151357.

6. Kupferschmidt K. Genome analyses help track coronavirus’ moves. Science. 2020;367(6483):1176–7. Epub 2020/03/14. doi: 10.1126/science.367.6483.1176. PubMed PMID: 32165562.

7. Holmes EC. The evolution and emergence of RNA viruses. Oxford; New York: Oxford University Press; 2009. xii, 254 p. p.

8. Hughes AL, Hughes MA. More effective purifying selection on RNA viruses than in DNA viruses. Gene. 2007;404(1-2):117–25. Epub 2007/10/12. doi: 10.1016/j.gene.2007.09.013. PubMed PMID: 17928171; PubMed Central PMCID: PMCPMC2756238.

9. Cagliani R, Forni D, Clerici M, Sironi M. Computational Inference of Selection Underlying the Evolution of the Novel Coronavirus, Severe Acute Respiratory Syndrome Coronavirus 2. J Virol. 2020;94(12). Epub 2020/04/03. doi: 10.1128/JVI.00411-20. PubMed PMID: 32238584.

10. Perutz MF, Kendrew JC, Watson HC. Structure and function of haemoglobin: II. Some relations between polypeptide chain configuration and amino acid sequence. Journal of Molecular Biology. 1965;13(3):669–78. doi: https://doi.org/10.1016/S0022-2836(65)80134-6.

11. Echave J, Spielman SJ, Wilke CO. Causes of evolutionary rate variation among protein sites. Nat Rev Genet. 2016;17(2):109–21. Epub 2016/01/20. doi: 10.1038/nrg.2015.18. PubMed PMID: 26781812; PubMed Central PMCID: PMCPMC4724262.

12. PDB. COVID-19/SARS-CoV-2 Resources 2020 [cited 2020 07/15/2020]. Available from: https://www.rcsb.org/news?year=2020&article=5e74d55d2d410731e9944f52.

13. Sedova M, Jaroszewski L, Alisoltani A, Godzik A. Coronavirus3D: 3D structural visualization of COVID-19 genomic divergence. Bioinformatics. 2020. Epub 2020/05/30. doi: 10.1093/bioinformatics/btaa550. PubMed PMID: 32470119; PubMed Central PMCID: PMCPMC7314196.

14. Goodsell DS, Zardecki C, Di Costanzo L, Duarte JM, Hudson BP, Persikova I, et al. RCSB Protein Data Bank: Enabling biomedical research and drug discovery. Protein Sci. 2020;29(1):52–65. Epub 2019/11/29. doi: 10.1002/pro.3730. PubMed PMID: 31531901; PubMed Central PMCID: PMCPMC6933845.

15. Porta-Pardo E, Godzik A. e-Driver: a novel method to identify protein regions driving cancer. Bioinformatics. 2014;30(21):3109–14. Epub 2014/07/30. doi: 10.1093/bioinformatics/btu499. PubMed PMID: 25064568; PubMed Central PMCID: PMCPMC4609017.

16. Yang Z, Bielawski JP. Statistical methods for detecting molecular adaptation. Trends Ecol Evol. 2000;15(12):496–503. Epub 2000/12/15. doi: 10.1016/s0169-5347(00)01994-7. PubMed PMID: 11114436; PubMed Central PMCID: PMCPMC7134603.

17. Denison MR, Graham RL, Donaldson EF, Eckerle LD, Baric RS. Coronaviruses: an RNA proofreading machine regulates replication fidelity and diversity. RNA Biol. 2011;8(2):270–9. Epub 2011/05/20. doi: 10.4161/rna.8.2.15013. PubMed PMID: 21593585; PubMed Central PMCID: PMCPMC3127101.

18. Wang C, Liu Z, Chen Z, Huang X, Xu M, He T, et al. The establishment of reference sequence for SARS-CoV-2 and variation analysis. J Med Virol. 2020;92(6):667–74. Epub 2020/03/14. doi: 10.1002/jmv.25762. PubMed PMID: 32167180; PubMed Central PMCID: PMCPMC7228400.

19. van Dorp L, Acman M, Richard D, Shaw LP, Ford CE, Ormond L, et al. Emergence of genomic diversity and recurrent mutations in SARS-CoV-2. Infect Genet Evol. 2020;83:104351. Epub 2020/05/11. doi: 10.1016/j.meegid.2020.104351. PubMed PMID: 32387564; PubMed Central PMCID: PMCPMC7199730.

20. Angelini MM, Akhlaghpour M, Neuman BW, Buchmeier MJ. Severe acute respiratory syndrome coronavirus nonstructural proteins 3, 4, and 6 induce double-membrane vesicles. mBio. 2013;4(4). Epub 2013/08/15. doi: 10.1128/mBio.00524-13. PubMed PMID: 23943763; PubMed Central PMCID: PMCPMC3747587.

21. Porta-Pardo E, Kamburov A, Tamborero D, Pons T, Grases D, Valencia A, et al. Comparison of algorithms for the detection of cancer drivers at subgene resolution. Nat Methods. 2017;14(8):782–8. Epub 2017/07/18. doi: 10.1038/nmeth.4364. PubMed PMID: 28714987; PubMed Central PMCID: PMCPMC5935266.

22. Tan J, Vonrhein C, Smart OS, Bricogne G, Bollati M, Kusov Y, et al. The SARS-unique domain (SUD) of SARS coronavirus contains two macrodomains that bind G-quadruplexes. PLoS Pathog. 2009;5(5):e1000428. Epub 2009/05/14. doi: 10.1371/journal.ppat.1000428. PubMed PMID: 19436709; PubMed Central PMCID: PMCPMC2674928.

23. von Brunn A, Teepe C, Simpson JC, Pepperkok R, Friedel CC, Zimmer R, et al. Analysis of intraviral protein-protein interactions of the SARS coronavirus ORFeome. PLoS One. 2007;2(5):e459. Epub 2007/05/24. doi: 10.1371/journal.pone.0000459. PubMed PMID: 17520018; PubMed Central PMCID: PMCPMC1868897.

24. Rogozin IB, Spiridonov AN, Sorokin AV, Wolf YI, Jordan IK, Tatusov RL, et al. Purifying and directional selection in overlapping prokaryotic genes. Trends Genet. 2002;18(5):228–32. Epub 2002/06/06. doi: 10.1016/s0168-9525(02)02649-5. PubMed PMID: 12047938.

25. Brown CJ, Johnson AK, Daughdrill GW. Comparing models of evolution for ordered and disordered proteins. Mol Biol Evol. 2010;27(3):609–21. Epub 2009/11/20. doi: 10.1093/molbev/msp277. PubMed PMID: 19923193; PubMed Central PMCID: PMCPMC2822292.

26. Peñarrubia L, Ruiz M, Porco R, Rao SN, Juanola-Falgarona M, Manissero D, et al. Multiple assays in a real-time RT-PCR SARS-CoV-2 panel can mitigate the risk of loss of sensitivity by new genomic variants during the COVID-19 outbreak. Int J Infect Dis. 2020;97:225–9. Epub 2020/06/12. doi: 10.1016/j.ijid.2020.06.027. PubMed PMID: 32535302; PubMed Central PMCID: PMCPMC7289722.

27. Álvarez-Díaz DA, Franco-Muñoz C, Laiton-Donato K, Usme-Ciro JA, Franco-Sierra ND, Flórez-Sánchez AC, et al. Molecular analysis of several in-house rRT-PCR protocols for SARS-CoV-2 detection in the context of genetic variability of the virus in Colombia. Infect Genet Evol. 2020;84:104390. Epub 2020/06/04. doi: 10.1016/j.meegid.2020.104390. PubMed PMID: 32505692; PubMed Central PMCID: PMCPMC7272177.

28. Klungthong C, Chinnawirotpisan P, Hussem K, Phonpakobsin T, Manasatienkij W, Ajariyakhajorn C, et al. The impact of primer and probe-template mismatches on the sensitivity of pandemic influenza A/H1N1/2009 virus detection by real-time RT-PCR. J Clin Virol. 2010;48(2):91–5. Epub 2010/04/21. doi: 10.1016/j.jcv.2010.03.012. PubMed PMID: 20413345.

29. Brault AC, Fang Y, Dannen M, Anishchenko M, Reisen WK. A naturally occurring mutation within the probe-binding region compromises a molecular-based West Nile virus surveillance assay for mosquito pools (Diptera: Culicidae). J Med Entomol. 2012;49(4):939–41. doi: 10.1603/me11287. PubMed PMID: 22897055; PubMed Central PMCID: PMCPMC3541937.

30. Ward JJ, Sodhi JS, McGuffin LJ, Buxton BF, Jones DT. Prediction and functional analysis of native disorder in proteins from the three kingdoms of life. J Mol Biol. 2004;337(3):635–45. Epub 2004/03/17. doi: 10.1016/j.jmb.2004.02.002. PubMed PMID: 15019783.

31. Cubuk J, Alston JJ, Incicco JJ, Singh S, Stuchell-Brereton MD, Ward MD, et al. The SARS-CoV-2 nucleocapsid protein is dynamic, disordered, and phase separates with RNA. bioRxiv. 2020. Epub 2020/06/18. doi: 10.1101/2020.06.17.158121. PubMed PMID: 32587966; PubMed Central PMCID: PMCPMC7310622.

32. Wang R, Hozumi Y, Yin C, Wei G-W. Mutations on COVID-19 diagnostic targets. arXiv. 2020. doi: arXiv:2005.02188.

33. Page AJ, Taylor B, Delaney AJ, Soares J, Seemann T, Keane JA, et al.: rapid efficient extraction of SNPs from multi-FASTA alignments. Microb Genom. 2016;2(4):e000056. Epub 2016/04/29. doi: 10.1099/mgen.0.000056. PubMed PMID: 28348851; PubMed Central PMCID: PMCPMC5320690.

34. Li H. A statistical framework for SNP calling, mutation discovery, association mapping and population genetical parameter estimation from sequencing data. Bioinformatics. 2011;27(21):2987–93. Epub 2011/09/08. doi: 10.1093/bioinformatics/btr509. PubMed PMID: 21903627; PubMed Central PMCID: PMCPMC3198575.

35. Webb B, Sali A. Protein Structure Modeling with MODELLER. Methods Mol Biol. 2017;1654:39–54. doi: 10.1007/978-1-4939-7231-9_4. PubMed PMID: 28986782.

36. Jaroszewski L, Li Z, Cai XH, Weber C, Godzik A. FFAS server: novel features and applications. Nucleic Acids Res. 2011;39(Web Server issue):W38–44. doi: 10.1093/nar/gkr441. PubMed PMID: 21715387; PubMed Central PMCID: PMCPMC3125803.

37. Fraczkiewicz R, Braun W. Exact and efficient analytical calculation of the accessible surface areas and their gradients for macromolecules. Journal of Computational Chemistry. 1998;19(3):319–33. doi: 10.1002/(sici)1096-987x(199802)19:3<319::Aid-jcc6>3.3.Co;2-3. PubMed PMID: WOS:000071747800006.

